# Structure based analysis of protein cluster size for super-resolution microscopy in the nervous system

**DOI:** 10.1101/845107

**Authors:** Chia-En Wong, Cheng-Che Lee, Kuen-Jer Tsai

## Abstract

To overcome the diffraction limit and resolve target structures in greater detail, far-field super-resolution techniques such as stochastic optical reconstruction microscopy (STORM) have been developed, and different STORM algorithms have been developed to deal with the various problems that arise. In particular, the effect of local structure is an important issue. For objects with closely correlated distributions, simple Gaussian-based localization algorithms often used in STORM imaging misinterpret overlapping point spread functions (PSFs) as one and this limits the ability of super-resolution imaging to resolve nanoscale local structures and leading to inaccurate length measurements. In the present study, we proposed a novel, structure-based, super-resolution image analysis method: structure-based analysis (SBA), which combines a structural function and a super-resolution localization algorithm. Using SBA, we estimated the size of fluorescent beads, inclusion proteins, and subtle synaptic structures in both wide-field and STORM images. The results showed that SBA has comparable and often superior performance to commonly used full-width-at-half-maximum parameters. We also demonstrated that SBA provides size estimations that corroborate previously published electron microscopy data.

## Introduction

Fluorescent microscopy is widely used in many biological fields to reveal molecular distributions and cellular structures. However, the resolution of optical microscopy is limited by the diffraction of light: two objects separated by a distance smaller than the diffraction limit cannot be distinguished separately ^1^. Under optical microscopy, the spatial distribution of photons from an isolated point source on the image plane is described in terms of the point spread function (PSF) of the microscope, which is generally expressed as the Airy pattern ^2^. In single molecule localization microscopy, this property can be exploited because the center of each isolated PSF can be estimated using localization algorithms, achieving spatial resolutions of a few nanometers ^3,4^. To resolve densely labeled objects separated by distances smaller than the diffraction limit, far-field, super-resolution techniques exploiting photoswitchable fluorophores have been developed. Examples include stochastic optical reconstruction microscopy (STORM) and photo-activated localization microscopy (PALM), in which different sets of fluorophores are activated, detected, and localized in different image frames ^5,6^. The temporal information registered in each frame of the series is then reconstructed into a pointillistic, super-resolution image. These techniques have been employed to study the nervous system and to resolve subtle neuronal structures such as synaptic protein localization, the presynaptic terminal, and the postsynaptic densities (PSDs), which cannot be resolved clearly using confocal microscopy ^7^.

In localization-based super-resolution imaging, a single emitting fluorophore is generally localized by fitting a two-dimensional Gaussian function with a constant background to the spatial distribution of the light intensity of each spot in the image series. The central position of this Gaussian function is then determined and recorded as the location of the fluorophore ^7^. Different localization algorithms can be used to reconstruct super-resolution images for different purposes ^8^. For example, fast-fitting algorithms, such as MUSICAL, have been proposed to decrease the time required for image reconstruction of single or multiple PSFs ^9^. Algorithms that process densely labeled samples, such as DAOSTORM, FALCON, and deconSTORM, have also been proposed ^10,11,12^. As a result, various localization algorithms with different PSF forms are now available to address different problems.

However, one important issue in localization microscopy is the effect of local structure, including that of objects with closely spaced fluorophores ^8,13,14^. Such structures are frequently encountered in studying the nervous system. For example, the PSD is a densely packed, disc-shaped protein cluster in the dendritic spines of the neurons ^15,16^. In many neurodegenerative diseases, abnormal proteins inclusion bodies—densely packed, spherical protein clusters are other examples ^17,18^. In such cases, PSFs generated by neighboring fluorophores overlap spatially. Because the localization process only records the central position of each fluorophore, and the information registered in the lateral width of the PSF is discarded during fitting, overlapping PSFs are misinterpreted as single ones. This limits the ability of pointillistic super-resolution imaging to resolve nanoscale local structures and to measure the length of objects with correlated distributions.

In the present work, we designed a novel, structure-based method to analyze the local dimensions of a localization cluster in super-resolution STORM images with better accuracy and precision, allowing the size of a localization cluster’s underlying structure to be estimated. Since previous studies using electron microscopy have found that, in cases of PSD and protein inclusion, the protein structures are disk-shaped and spherical, respectively ^16,17^. We incorporated this structural information of the underlying protein structures to better analyze the physical size of the cluster structure. In subsequent studies, we proposed a novel STORM image analysis method that uses structure-based fitting algorithms, naming the method “structure-based analysis (SBA).” The technique estimates, using predetermined structural information as mentioned, the size parameters of a localization cluster with a corresponding structural function. Using synthetic data, we then tested SBA with spherical structural function and validated it by applying it to standardized, 100-nm fluorescent microspheres. The method showed superior performance compared to the commonly used full-width-at-half-maximum (FWHM) parameter. Furthermore, we analyzed two relevant cluster-forming proteins of the nervous system—TAR DNA-binding protein 43 (TDP-43) clusters and the PSD marker, PSD-95. In this way, we demonstrated that SBA displayed better performance than commonly used parameters, as well as comparable results to previously published electron microscope data.

## Materials and methods

### Fluorescent microsphere sample preparation

To prepare the fluorescent bead samples, 100-nm TetraSpeck microspheres (Invitrogen) were deposited onto 20-mm glass coverslips, which were then placed at the bottom of 12-well culture plates. Next, 100 μL of a 1:150 microspheres dilution was added to each well. The coverslips, with loaded microspheres, were kept at 4°C overnight, and 500 μL distilled water was then added. The microsphere samples were kept at 4°C for storage.

### HEK 293T cell culture

Human epithelial kidney (HEK 293T) cells were maintained in Dulbecco’s Modified Eagle’s Medium (Gibco) supplemented with 10% fetal bovine serum (Invitrogen, San Diego, CA) at 37°C in an atmosphere of 95% air and 5% CO_2_.

### Sample preparation of TDP-43 protein inclusions

Before the TDP-43 inclusion bodies were prepared, 20-mm glass coverslips were coated with 1 μg/mL of laminin at 37°C for 4 hours and with 1 mg/mL of poly-D-lysine at room temperature for 20 minutes. The coated coverslips were then placed in the bottom of 12-well cell culture plate, and HEK 293T cells were seeded onto the culture plate at a density of 2 × 10^5^ per well. Forty-eight hours after seeding, the HEK 293T cells were transfected with plasmid pMM403-TDP43-GFP to over-express TDP-43; 36 hours after transfection, the transfected HEK 293T cells were treated with MG132 (Sigma-Aldrich) at a concentration of 20 μM for 16 hours. Next, the prepared HEK 293T cells were fixed with paraformaldehyde (PFA) for 20 minutes. Excess PFA was washed away using phosphate buffered saline (PBS) three times, and the TDP-43 protein inclusion cell samples were stored in PBS at 4 °C.

### Establishment and maintenance of cortical primary neuron culture and PSD-95 sample preparation

All animal experimental procedures were approved by the Institutional Animal Care and Use Committee at the National Cheng Kung University, in accordance with National Institute of Health guidelines. The sample of PSD-95 clusters were prepared from primary mouse cortical neurons. Before culture, 20-mm glass coverslips were coated with 1 mg/mL of poly-D-lysine at room temperature for 20 minutes; the coated coverslips were then placed into the bottom of 12-well culture plates. Next, mouse pups at postnatal day 0 were sacrificed and their cerebral cortices excised under a dissection microscope. Cortical tissues were triturated for disaggregation and plated on poly-D-lysine-coated glass coverslips in Neurobasal medium containing B27 serum-free supplement and antibiotics (50 U/mL penicillin and 50 μg/mL streptomycin). Cultures were incubated at 37°C in an atmosphere of 95% air, 5% CO_2_, and 90% relative humidity. Half of the growth medium was replaced every 2–3 days. The primary cultured neurons were maintained for 19 days prior to immunostaining.

### Immunofluorescence

To apply immunofluorescence, the prepared cell cultures were fixed using 4% PFA for 20 minutes at room temperature. The cells were then washed in PBS for 3 x 10 minutes and blocked using 5% bovine serum albumin in phosphate buffered saline with tween 20 and 0.5% Triton-X. After blocking, the cells were hybridized using the primary antibodies (anti-TDP43, ProteinTech, anti-PSD-95; Abcam) overnight at 4°C. After washing in PBS for 3 x 10 minutes, the samples were incubated in the secondary antibody (Alexa Fluor 647; Life Technologies) for 1 hour at room temperature.

### Imaging buffer preparation

We used the imaging buffer composition suggested in a previously reported protocol ^31^. Buffer A was composed of 10 mM TRIS (pH 8.0) and 50 mM NaCl, while buffer B was composed of 50 mM TRIS (pH 8.0), 10 mM NaCl, and 10% glucose. GLOX solution (1 mL) was prepared by vortex mixing a solution of 56 mg glucose oxidase, 200 μL catalase (17 mg/mL), and 800 μL of buffer A. MEA solution (1 M, 1 mL) was prepared using 77 mg of MEA and 1 mL of 0.25 N HCl. In each chamber of the 20-mm glass coverslips, 500 μL of imaging buffer was prepared by mixing 5 μL of GLOX solution, 50 μL of MEA solution, and 445 μL of buffer B on ice. This imaging buffer was used as the medium for the TDP-43 and PSD-95 samples. After the chamber was filled, it was covered immediately to avoid replenishing of the dissolved oxygen.

### STORM imaging

The microscope was constructed around an Olympus IX-83 automated inverted microscope. For illumination, an objective-type, total internal reflection fluorescence, oil-immersion objective (APON 60×; NA 1.49 total internal reflection fluorescence; Olympus) was used. A multiline laser source (405, 488, 561, and 640 nm; Andor Technology) was used for excitation and activation. Single-molecular localization signals were separated using appropriate filters (Andor Technology) and detected using electron-multiplying charge-coupled device camera (iXon Ultra 897; Andor Technology). Before dynamic image movie acquisition, conventional fluorescence images were acquired to determine the region of interest. Specifically, 10,000 frame image series were recorded with an exposure time of 50 ms (20 frames per second). The acquired image series were then analyzed using MetaMorph® Super-Resolution System (Molecular Devices) to generate reconstructed STORM images.

### Numerical fitting of SBA analysis in STORM images

SBA image analysis was performed on the localization number line profiles acquired from the STORM images and measured in MetaMorph® (Molecular Devices) software. The derivation of numerical fitting functions is described in Supplementary Note 1 and 2. The fitting was performed using the MATLAB software curve fitting toolbox (cftool), and the custom equation mode was used with the numerical SBA fitting function.

## Results

### Structure-based analysis

The idea of SBA arose because we wished to carry out super-resolution localization microscopy on many biological targets with known structures that had already been well-studied under electron microscopy^16,18^ but lacked optimal parameters to describe their size. Conventional estimation parameters used in diffraction-limited imaging techniques, such as FWHM, are still used in localization microscopy ^19,20^. However, these may be suboptimal because the total widths of localization clusters in super-resolution images do not account for the lateral width of the PSFs, which are reduced in localization process.

SBA overcomes this issue by adopting previously determined structural information. That is, specific parameters describing structure size are incorporated into the estimation algorithm in the form of structural function. In the present study, we briefly discussed the SBA algorithm.

The spatial probability distribution of the photons emitted from a single emitting fluorophore adhering to a surface can be described by the Poisson distribution ^21^, which can be mathematically modeled in terms of Gaussian approximation function denoted by *G*(*x*), whereby “x” is the distance from the center of the distribution— that is, from the emitting fluorophore ^22^.

The spatial emitting profile of a fluorescence-labeled structure emitting homogeneously distributed fluorophores from its surface, denoted by *D*(*x*), is the convolution of the function of a specific structure *S*(*x*) and the distribution profile of a single emitting fluorophore, as follows: 

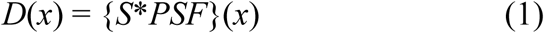

The function *D*(*x*) describes the total spatial distribution profile of the photons emitted from the structure *S*(*x*). Under STORM microscopy, the localization process records the central position of each PSF of randomly emitting fluorophores, so processed STORM images represent the total spatial distribution profile of the emitted photons captured and localized in every image frame, which is equivalent to the structure-based function *D*(*x*). SBA exploits this property to analyze the size of the underlying structure by least-square fitting of equation 1 to the line localization profile obtained from the STORM image. The explicit derivation is given in Supplementary Note 1, accompanied by a detailed discussion.

### SBA of a spherical structural function

We characterized an SBA fitting algorithm with a spherical structural function and compared the results of SBA with those of FWHM. Synthetic examples of two cylinders with diameters of 200 nm and 400 nm were used (**Fig. 1a**), and the cross-sectional line profile of the cylinders is shown in **Figure 1b**. The diameters of the cylinders were poorly resolved in diffraction-limited wild-field images (**Fig 1c**). STORM images of the cylinders were generated as described in the Materials and Methods section, and the cross-sectional line profiles along the dashed lines in **Figure 1d** of the STORM images were analyzed by SBA with a spherical structural function. A detailed derivation of SBA with a spherical structural function is given in Supplementary Note 2.

**Figure 1.**
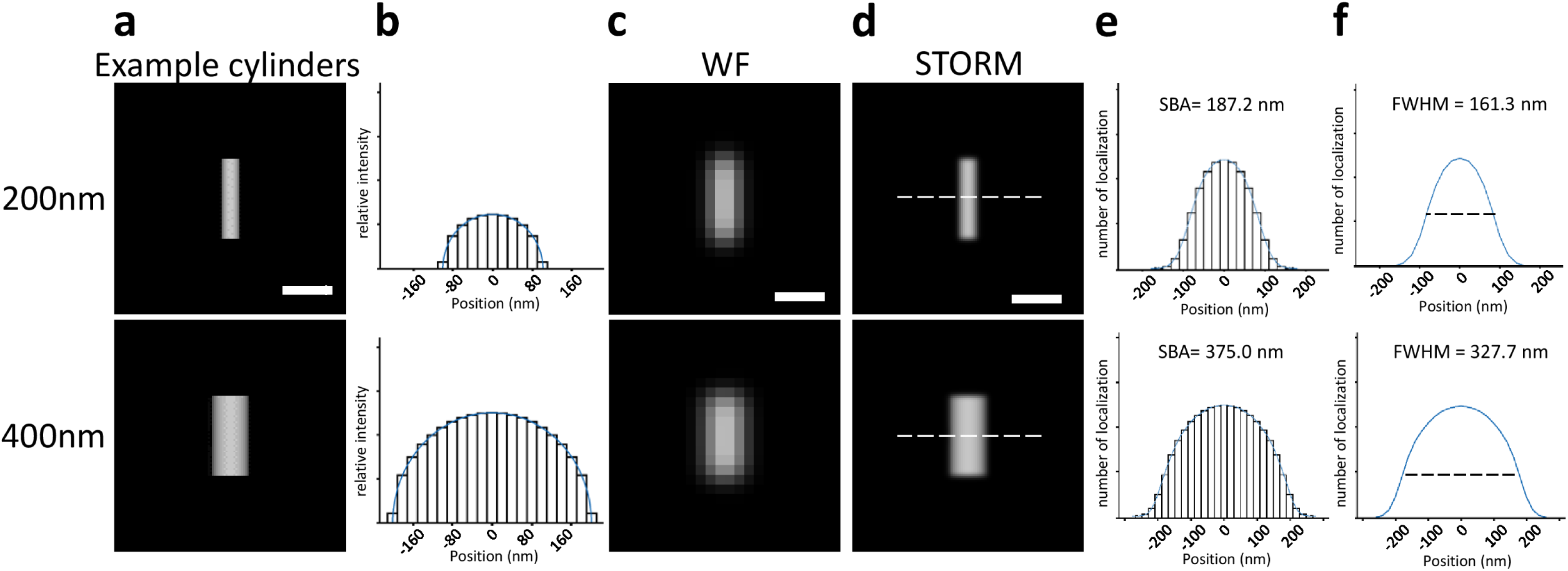
Demonstration of SBA with spherical function on synthetic cylindrical samples. (**a**) Synthetic examples of two cylinders with diameters of 200 nm and 400 nm. Scale bar: 500 nm. (**b**) Cross-sectional line profile of synthetic cylinders with diameters of 200 nm and 400 nm, respectively (upper panel: 200 nm, lower panel: 400 nm) (**c and d**) Representative wild-field or STORM images of the cylinders (upper panel: 200 nm, lower panel: 400 nm). Scale bar: 500 nm. (**e**) Demonstration of SBA of the cylinders. Line profiles were obtained along the dashed line, as shown in the STORM images (upper panel: 200 nm, lower panel: 400 nm). (**f**) FWHMs in the STORM images of the synthetic cylinders with diameters of 200 nm (upper panel) and 400 nm (lower panel).

The SBA-estimated diameters were 187.2 nm and 375.0 nm for the two cylindrical emitters with diameters of 200 nm and 400 nm, respectively (**Fig. 1e**). For comparison, the FWHMs of the cross-sectional line profiles along the dashed lines in **Figure 1d** were also measured; the measured FWHMs were 161.3 nm and 327.7 nm, respectively (**Fig. 1f**). Thus, SBA was superior to FWHM in its ability to estimate the size of specific structures.

### SBA diameter estimation using 100-nm fluorescent microspheres

To further test the performance of SBA under realistic conditions, 100-nm fluorescent microspheres were used as standard targets for wild-field images (**Fig. 2a**). We fixed 100-nm fluorescent beads onto glass coverslips and photographed them at 20 Hz to generate 10,000-frame image stacks. STORM images were then reconstructed from the image stacks (**Fig. 2b**). Line profiles were obtained along the principle axis of the spheres, which was the long axis of the cluster indicated by the dashed line in **Figure 2b**. SBA with the spherical structural function was used to estimate the size of the fluorescent microspheres from the STORM images (**Fig. 2c**). The results showed an average diameter of 98.2 ± 9.2 nm (mean ± standard deviation [SD], n = 123).

**Figure 2.**
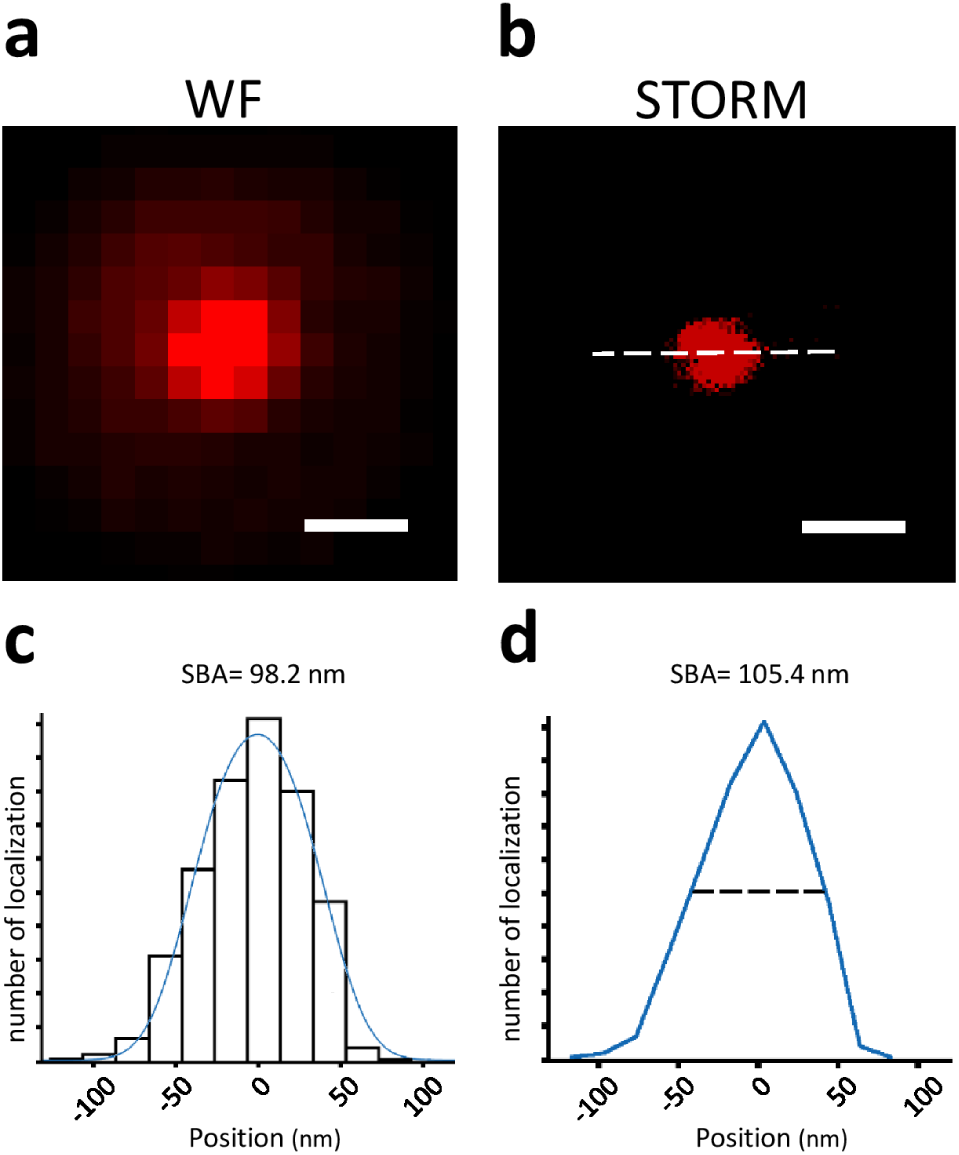
SBA of 100-nm fluorescence microspheres. (**a and b**) Wild-field and STORM images of a 100-nm fluorescent microsphere. Scale bar: 300 nm. (**c**) SBA performed on the STORM image of the microsphere. The line profile was obtained along the dashed line shown in **b**. (**d**) FWHM of the microsphere, the line profile was obtained along the dashed line shown in **b**.

We compared the results of SBA to those of the corresponding FWHMs of the microspheres in the STORM images; the average FWHM of the fluorescent microspheres in the STORM images was 105.4 ± 50.9 nm (mean ± SD; n = 123; **Fig. 2d**). The comparison showed that both SBFA and FWHM yielded a close approximation to the true size of the fluorescent beads, with SBA being slightly more accurate. However, the standard deviation of microsphere diameters estimated by SBA was 9.2 nm, while the standard deviation of the corresponding FWHMs was 50.9 nm, indicating that SBA yielded less dispersion than FWHM. Taken together, these results demonstrated SBA is precise in its estimation of diameter, and that it is comparable or even superior to FWHM.

### SBA in mammalian cells: Analysis of TDP-43 cluster size

Next, the neurobiological applications of SBA were demonstrated by estimating TDP-43 protein cluster sizes. TDP-43 is a multifunctional DNA/RNA binding protein with an important role in neurodegenerative diseases such as amyotrophic lateral sclerosis (ALS) and frontotemporal lobar degeneration (FTLD) ^23^. Past studies have identified TDP-43 inclusion bodies as the pathological hallmark of ALS/FTLD and proposed that the clustering and self-aggregation of TDP-43 is involved in the pathogenesis of these neurodegenerative diseases ^24,25^.

In the present study, we used SBA to study the cluster size of TDP-43. The samples of TDP-43 protein clusters were prepared as described in the Materials and methods section. Briefly, TDP-43 was transfected by lipofectamine and overexpressed in an HEK 293T cell line. After transfection for 24 hours, the protease inhibitor MG-132 was added to the cell culture to allow expression and formation of TDP-43 clusters. The TDP-43 molecules were photographed, and image stacks containing 10,000 frames were captured at 20 Hz. TDP-43 protein clusters were identified as puncta structures in both wild-field and STORM images reconstructed from the image stacks, as shown in **Figures 3a and 3b**. In the STORM images, the line profiles were obtained along the principal axes, which were defined as the long axes of the clusters (**Fig. 3c, 3d, and 3e**). The SBA of the TDP-43 clusters in regions A to C are displayed in the lower panels. The SBA-estimated diameters of the TDP-43 clusters ranged from 50 to 550 nm (332.4 ± 137.7 nm; mean ± SD; n = 59), which is comparable with previously reported sizes of TDP-43 oligomers and inclusions measured by electron microscopy in previous studies ^26,27^. The SBA results of the TDP-43 molecules demonstrated that SBA can be applied to neurobiologically relevant targets and that it can estimate the size of protein clusters in STORM images.

**Figure 3.**
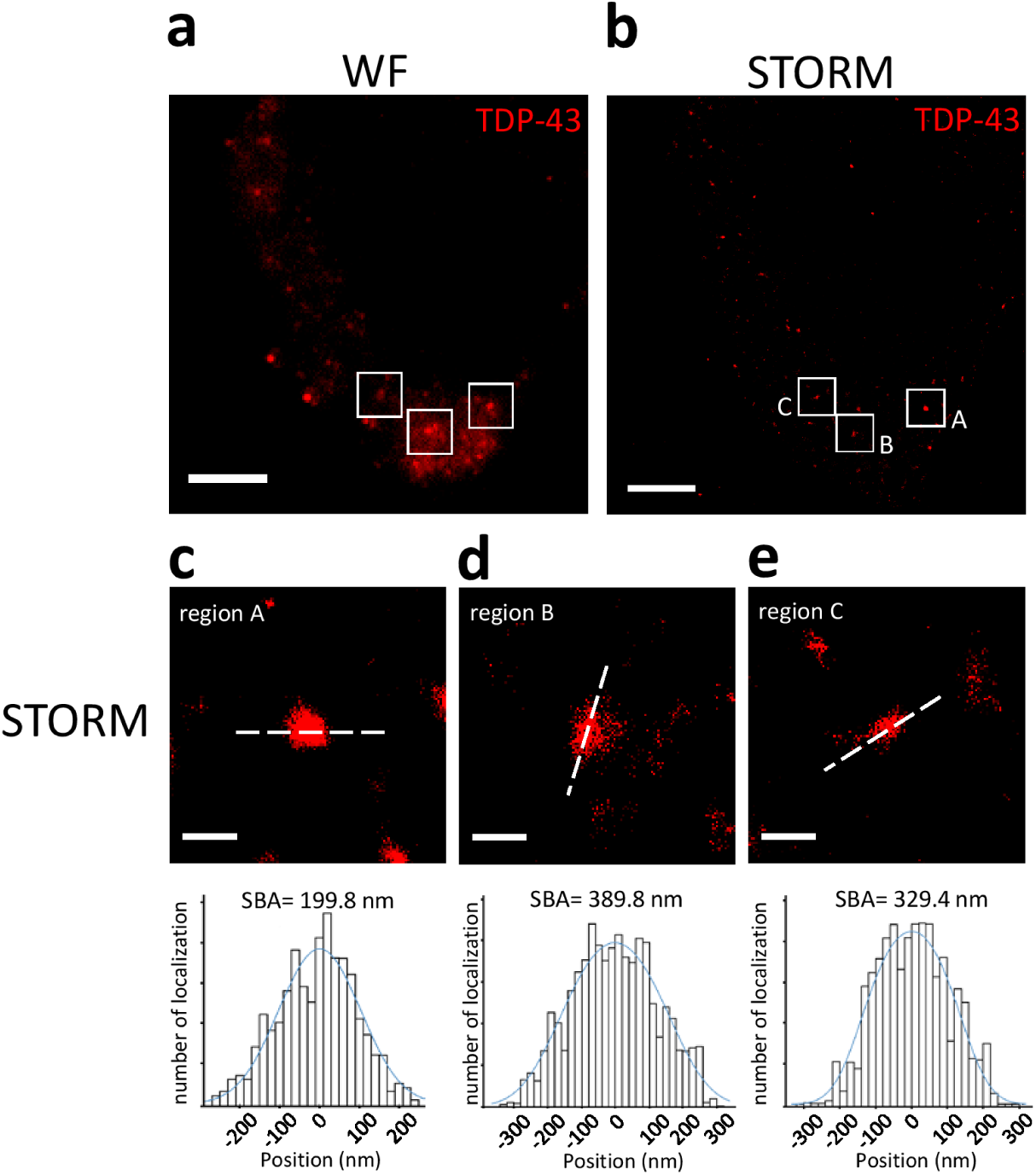
Application of SBA in TDP-43 cluster size estimation. (**a and b**) Wild-field and STORM images of TDP-43 protein clusters in HEK293T cells. Scale bar: 5 μm. (**c, d, and e**) Representative magnification images showing a single TDP-43 cluster in region A-C of the STORM images (upper panels). Scale bar: 500 nm. The lower panels display the SBA of the TDP-43 clusters. The line profiles were obtained along the principle axes shown as the dashed lines in the upper panels.

### Application of SBA in the neuronal system: Analysis of PSD size in neuronal cells

We next applied SBA in a neurobiological setting to estimate the size of the PSDs, which are disc-shaped protein clusters within the dendritic spines of neuronal cells, with sizes ranging from 300–700 nm^16,28^. To study PSD size, primary mouse cortical neuronal cultures were prepared on glass coverslips using a previously described method with some minor modifications. The cultured neurons were maintained for 19 days for maturation, and the PSD structural protein, PSD-95, was imaged using both wild-field immunofluorescence and STORM. We observed PSD-95 clusters forming puncta structures and distributing along the dendrites of the neurons in both the wild-field and STORM images (**Fig. 4a and b**). The average FWHM of the PSD-95 puncta in the wild-field images was 824.1 ± 260.3 nm (mean ± SD; n = 71) (**Fig 4c**). From the STORM images, the long axis of each PSD-95 puncta was identified as the principal axis of the disc-shaped structure corresponding to the diameter of the disc-shaped PSD-95 clusters. The line profiles were acquired along the principle axes and analyzed by SBA (**Fig 4d, right**). The average diameter of the PSD-95 disc-shaped cluster was 285.2 ± 78.0 nm (mean ± SD; n = 71). Next, we compared the SBA results of the PSD-95 clusters in the STORM images to the FWHM results. The FWHM estimation of the mean PSD-95 size in STORM images was 192.9 ± 58.9 nm (mean ± SD; n = 71) (**Fig 4d, left**), which was smaller than the SBA estimated diameter. Importantly, the result of SBA estimated PSD cluster size was comparable with previously published PSD diameters ranging from 250 nm to 450 nm reported by previous studies using electron microscopy^28,29,30^.

**Figure 4.**
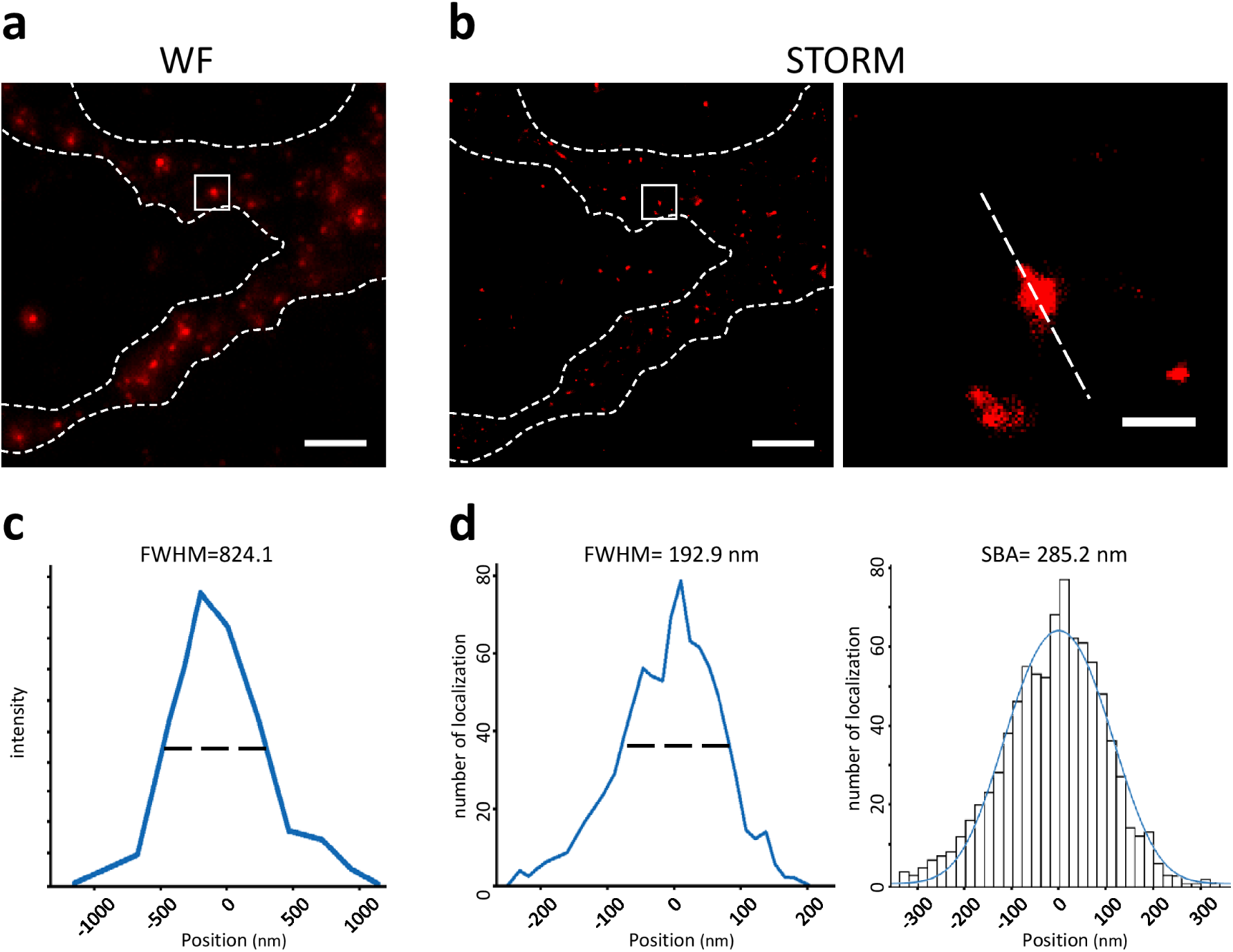
Application of SBA in PSD-95 length estimation. (**a**) Wild-field image of PSD-95 protein clusters in dendrites of primary mouse cortical neurons. Scale bar: 5 μm. **(b)** STORM images of PSD-95 protein clusters in dendrites of primary mouse cortical neurons. Scale bar: 5 μm (500 nm in magnified region). (**c**) FWHM of the PSD-95 cluster from wild-field image shown in **a**. (**d**) FWHM and SBA of the PSD-95 cluster. The line profile was obtained along the principle axis, shown as the dashed lines in **c**.

Taken together, these results demonstrated that SBA can estimate the sizes of subtle neuronal structures, such as PSDs. The SBA estimation of PSD-95 diameter was comparable with that of previously published electron microscope studies and suggested that SBA provided superior estimation parameters than FWHM.

## Discussion

In the present study, we described SBA, a cluster size estimation method for super-resolution STORM microscopy. By incorporating the structural function corresponding to the underlying protein structure of a localization cluster, SBA overcame the limitations posed by local structures, in which misinterpreted overlapping PSFs limit the ability of super-resolution imaging to measure the length of objects with correlated distributions^8,13,14^.

SBA was verified with both synthetic cylindrical emitters and fluorescent microspheres. The results demonstrated that SBA closely estimated the size parameter of specific structures and showed superior performance to FWHM.

Next, we applied SBA to cells overexpressing TDP-43. The SBA-estimated TDP-43 cluster sizes were comparable with previously reported size ranges of TDP-43 oligomers and inclusions measured using electron microscopy^26,27^. The different sizes observed here might represent different growth stages of TDP-43 inclusions, and the results showed the ability of SBA to quantify protein sizes under far-field super-resolution microscopy.

Furthermore, SBA was applied in a neurobiological context to estimate the size of the PSDs. The results showed superior performance of SBA to FWHM to estimate subtle PSD structures that were unresolvable in wild-field images, suggesting that it could analyze the dimensions of PSDs by providing quantitative information, corroborating previous electron microscopy studies^28,29,30^.

Although some previous studies have addressed the issue of local structure in localization-based, far-field super-resolution microscopy, these methods required modifications to the localization algorithm ^9,10,11,12^. In contrast, SBA served as a post-processing method that required no modifications to the localization process and could estimate the size of localization clusters in ways that other post-processing software could not ^32^.

We demonstrated that SBA can estimate molecular cluster sizes in far-field, super-resolution STORM images, and that SBA was comparable and often superior to FWHM. We also found that SBA yields size estimations comparable to previously published electron microscopy data.

SBA will be particularly useful in studying the size of molecular clusters with known or assumed shapes, such as protein inclusions, membrane rafts, and neuronal processes. We believe that SBA is a step towards more precise size estimation in fluorescence microscopy.

## Compliance with ethical standards

The authors declare no competing financial and non-financial interests.

## Author contributions

C.E.W. and K.J.T. designed the experiments. C.E.W. derived the equations and performed the experiments. C.E.W., C.C.L and K.J.T. interpreted the results. C.E.W, C.C.L and K.J.T. wrote the manuscript. All authors contributed to editing the manuscript.

## Acknowledgements

This study was performed, in part, with support from the Ministry of Science and Technology of Taiwan (MOST-105-2628-B-006-016-MY3 and MOST-106-2628-B-006-001-MY4).

